# Enhanced fluorescence from semiconductor quantum dot-labelled cells excited at 280 nm

**DOI:** 10.1101/2021.11.08.467709

**Authors:** Mollie McFarlane, Nicholas Hall, Gail McConnell

## Abstract

Semiconductor quantum dots (QDs) have significant advantages over more traditional fluorophores used in fluorescence microscopy including reduced photobleaching, long-term photostability and high quantum yields, but due to limitations in light sources and optics, are often excited far from their optimum excitation wavelengths in the deep-UV. Here, we present a quantitative comparison of the excitation of semiconductor QDs at a wavelength of 280 nm, compared to the longer wavelength of 365 nm, within a cellular environment. We report increased fluorescence intensity and enhanced image quality when using 280 nm excitation compared to 365 nm excitation for cell imaging across multiple datasets, with a highest average fluorescence intensity increase of 3.59-fold. We also find no significant increase in photobleaching of QDs associated with 280 nm excitation.

## 1. Introduction

Semiconductor quantum dots (QDs) have several significant advantages over traditional fluorophores commonly used in fluorescence microscopy which make them appealing for cell imaging applications. QDs exhibit high fluorescence efficiency, including high quantum yields and high molar extinction coefficients [1] and a strong resistance to photobleaching, with a photostability around 1000 times greater than conventional dyes [1, 2, 3]. The fluorescence emission wavelength of QDs depends on the size of the dot, allowing for the excitation of several QD colours with a single wavelength which offers advantages for multiplex imaging.

QDs are usually excited in the blue or near-UV region of the spectrum due to availability of suitable light sources, but the theoretically optimum excitation wavelength, given by their excitation spectra, lie in the deep-UV. Recent innovations in light emitting diode (LED) technology have produced high-brightness deep-UV emitters with optical powers in the 100 mW range [4]. LEDs with wavelengths of 280nm not only report some of the highest quantum efficiencies in the deep-UV [4] but are of particular interest in microscopy due to both the possible resolution enhancement associated with detecting at this wavelength [5] and their overlap with the excitation spectrum of many fluorophores. 280 nm light has previously been used to image cellular autofluorescence by exploiting endogenous fluorophores such as the aromatic amino acid tryptophan [5]. More recently, Microscopy with Ultraviolet Surface Excitation (MUSE) [6, 7, 8, 9] has identified that 280 nm light can be applied to excite fluorescence from different conventional fluorescent dyes whilst reporting further advantages of 280 nm excitation in tissue histology such as improved image contrast due to surface-limited excitation. A further application of MUSE has identified the suitability of QDs for excitation at 280 nm and applied this to multiplexed protein-specific imaging [10]. However, this work did not provide a quantitative insight into the benefits of excitation at this lower wavelength.

Based on the absorption spectrum, we can hypothesise that 280 nm excitation of semiconductor QDs will lead to an increase in fluorescence intensity. However, due to the complex nature of biological systems, it is not guaranteed that this will translate to in-vitro experimental conditions. For example, previous reports have found that the fluorescence of semiconductor QDs can be quenched by various chemical compounds existing within the cellular environment such as nucleotides and amino acids [11]. Further to this, it has been reported that semiconductor QDs can be quenched in the presence of bovine serum albumin (BSA) [12, 13], a key component to any immunolabelling protocol.

In this study, we label mammalian cells with commercially available QDs and provide a quantitative analysis of the fluorescence intensity of QDs excited with 280 nm light and 365 nm light, a wavelength already routinely used in light microscopy for the excitation of common fluorophores. We quantify the increase in fluorescence intensity of QDs within the cell, using two sizes of QDs, although we expect the proposed increase in fluorescence intensity to apply to all emission varieties of commercial semiconductor QDs due to their common absorption spectra. We also determine whether the increased energy associated with 280 nm excitation affects the photobleaching rate of QDs.

## 2. Materials and Methods

### 2.1. Chemicals and Reagents

All chemicals and reagents were purchased from ThermoFisher Scientific or Sigma Aldrich. Commercial CdSe/ZnS QD Streptavidin conjugates were purchased from ThermoFisher Scientific.

### 2.2. Microscope Set-Up

Imaging was performed on a modified Olympus BX50 microscope. One significant obstacle in using 280 nm light in a commercial microscope is low transmission of 280 nm light through most types of glass. This makes imaging in epifluorescence mode difficult, requiring the use of rare and costly quartz objective lenses. To overcome this, the specimen was instead illuminated from below the stage, in transmission mode. To achieve this, the condenser unit was removed from the microscope and quartz optics were installed along the optical bench to guide illumination light from the 280 nm LED to the specimen plane (Figure 1). The 280 nm LED was provided by CoolLED Ltd (LG Innotek LEUVA66H70HF00), with an optical power of 100 mW, a Lambertian emission profile and a peak wavelength of 278 nm with a FWHM of 10 nm. Due to the heavily patterned nature of the chip surface, critical illumination, which focuses an image of the chip onto the specimen, is unsatisfactory, particularly when performing quantitative fluorescence intensity analysis. For this reason, illumination optics were designed to provide as homogenous illumination as possible to the specimen by ensuring that the illumination light was not at a focal point when reaching the specimen. Illumination light was collimated by a set of 2 plano convex lenses (Edmund Optics 49-965, f = 30 mm), placed together to increase the collection efficiency of 280 nm photons, through a 300/50 nm excitation filter (Edmund Optics 12-093). Light was then relayed through two quartz lenses (Thorlabs LA4380-UV f = 100 mm, LA4148-UV, f = 50 mm.) The spot size of the illumination light was then reduced using a quartz lens pair (Thorlabs LA4148-UV f = 50 mm, LA4052-UV, f = 30 mm) to roughly match the field of view of the microscope and reflected to the specimen plane at 90° using a UV-enhanced aluminium mirror (Thorlabs PF10-03-F01). The 365 nm excitation was achieved in epifluorescence mode by attaching a CoolLED pE300white SB illuminator system to the epifluorescence port of the BX50 microscope and aligning for Köhler illumination. A Teledyne Photometrics CoolSnap HQ2 camera was used as a detector. Images of cells were acquired with a 10X/0.4NA objective (Olympus UPLXAPO10X) lens. All acquisitions were performed using *µ*Manager [14]. To measure the illumination uniformity, a fluorescent microscope slide (Chroma 92001) was used as a specimen. Images of the fluorescent slide were acquired with both 365 nm and 280 nm illumination.

**Figure 1:**
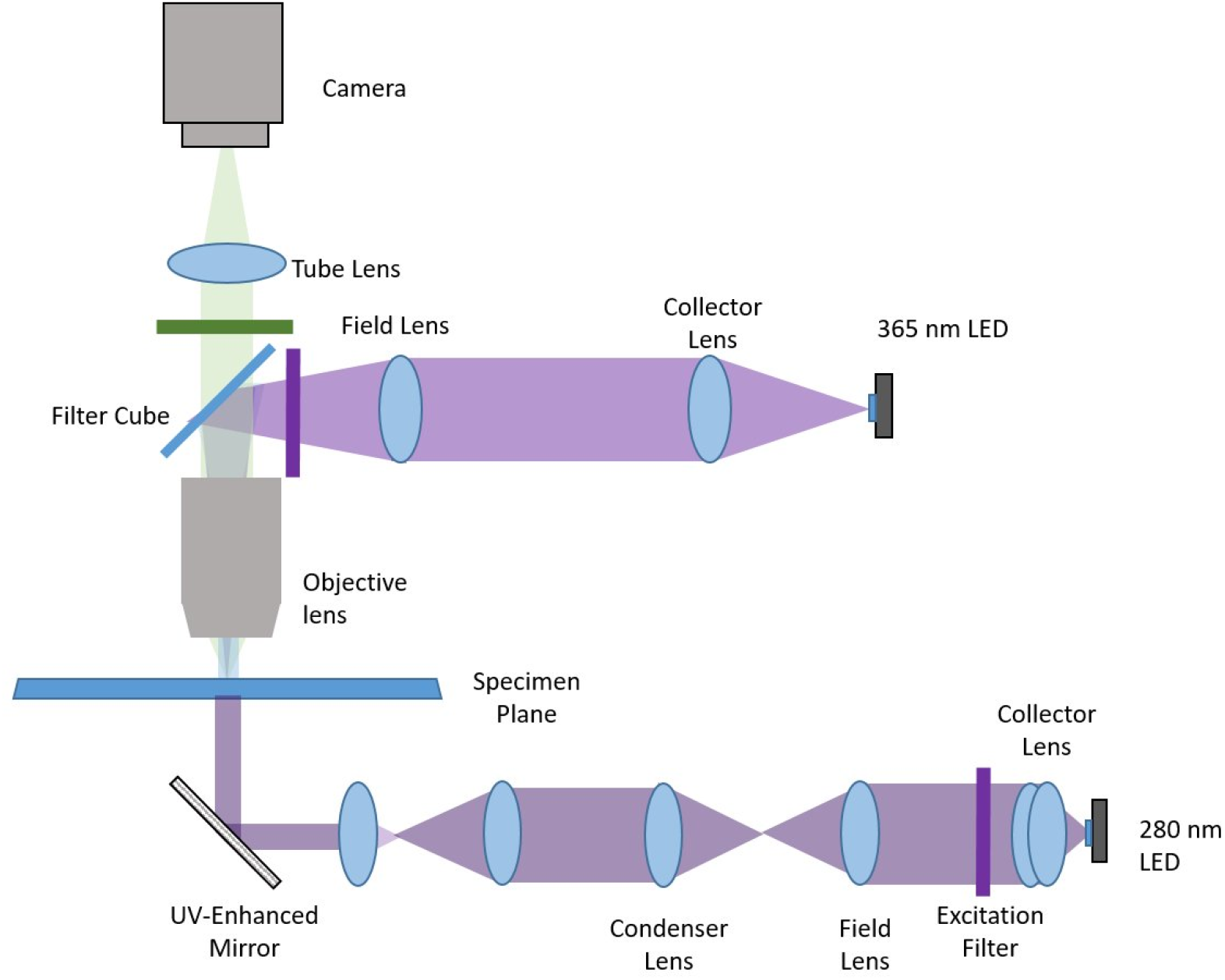
An Olympus BX50 microscope was modified to deliver 365 nm light and 280 nm light to the specimen plane. 365 nm light was delivered to the specimen through the epifluorescence pathway of the commercial microscope. Due to low transmission of 280 nm light through most glass, 280 nm light was delivered to the specimen in transmission fluorescence using quartz optics. Light from the 280 nm LED was collected using two quartz plano-convex lenses to ensure maximum collection efficiency and relayed to the specimen using quartz lenses to achieve the best possible illumination uniformity. A pair of quartz plano-convex lenses were used to narrow the collimated light to a spot size that roughly matched the field of view of the microscope objective. A UV-enhanced aluminium mirror was placed at a 90° angle below the stage to reflect the 280 nm light to the specimen. Images were collected using a 10x/0.4 NA microscope objective and relayed to the detector through a filter cube.

### 2.3. Cell Preparation and Immunolabelling with Commercial QDs

HeLa cells were grown on quartz coverslips (Alfa Aesar 43211) coated in fibronectin and cultured in Dulbecco’s Modified Eagle Medium (DMEM) until a confluency of around 50% was reached. Antibody labelling was performed as follows with three 5 minute washes in PBS between subsequent steps. First, cells were fixed with 4% formaldehyde for 20 minutes and permeabilized with 0.25% Triton X-100 in PBS for 15 minutes. Next, blocking of endogenous biotin was performed using a commercial blocking kit (Invitrogen, E21390). Cells were incubated in component A for 45 minutes, followed by component B for 45 minutes. Further blocking was performed using 6% BSA in PBS for 1 hour to prevent nonspecific binding of antibodies. An anti-alpha-tubulin monoclonal antibody (Sigma Aldrich T6199) was chosen as the primary antibody. Cells were incubated in this antibody at a dilution of 1:200 overnight. After incubation with primary antibody, the cells were incubated with a biotinylated secondary antibody (ThermoFisher Scientific 13-4013-85) at a dilution of 1:250 for two hours. Two wavelengths of QD conjugates were chosen from a QD-Streptavidin Sampler Kit (Thermofisher Scientific Q10151MP), one emitting at 525 nm (QD525) and one emitting at 605 nm (QD605), in order to compare the increased fluorescence from multiple QD sizes. Cells were treated with a 20 nM concentration of QD525 or a 40 nM concentration of QD605 for two hours. Cell-coated coverslips were subsequently mounted on a quartz microscope slide (Alfa Aesar 42297) using a gelvatol mounting medium.

To prepare unlabelled cells as a control sample, HeLa cells were grown on quartz coverslips as described previously and fixed with 4% formaldehyde. Cell-coated coverslips were then mounted on a quartz microscope slide using a gelvatol mounting medium.

### 2.4. Cell Imaging

Cell imaging was performed using the previously described microscope shown in Figure 1. The power of excitation light at the specimen plane was measured using a Thorlabs power meter with a UV-extended photodiode sensor (PM100A, S120VC). When measuring each wavelength of light, the detection wavelength was programmed into the power meter to account for wavelength dependency in the detector. LED drive currents were adjusted to ensure the optical power at the specimen plane was equal for each wavelength of light.

Images of a blank quartz coverslip-slide combination were obtained at the same LED power and camera exposure as QD images to measure the background of each image.

The autofluorescence of unlabelled HeLa cells was measured to ensure any increase in fluorescence in QD labelled cells was due to increased QD excitation efficiency rather than increased cellular autofluorescence. Unlabelled HeLa cells were imaged with the same camera exposure and LED power as the QD image pair. The mean autofluorescence intensity associated with each wavelength of light was subsequently subtracted from mean QD intensities during data analysis.

For imaging of QD525 labelled cells, a 525/50 bandpass emission filter was used (Semrock FF03-525/50-25). For imaging of QD605 labelled cells, a 561 LP emission filter was used (Semrock BLP02-561R-25). Cells were imaged first with 365 nm excitation light, then the same region was immediately imaged with 280 nm excitation of the same optical power. The power at the specimen plane and camera exposure were set to 3.8 mW and 500 ms, respectively, for QD525-labelled cells, and 4.8 mW and 100 ms for QD605-labelled cells. Optical powers and camera exposure times were chosen to avoid overexposure in images due to differing concentrations and quantum yields between QD525 and QD605 samples. To compare photobleaching of QDs illuminated with different excitation wavelengths, QD605-labelled cells were exposed to each wavelength of light for 8 hour periods. An optical power of 0.45 mW at the specimen plane was chosen for both wavelengths of light to avoid degradation of the gelvatol mountant over long periods of irradiation with high-intensity 280 nm light. Cells were irradiated constantly with light and imaged once every 10 minutes at a camera exposure of 500 ms. Experiments were repeated in triplicate.

### 2.5. Data Analysis

The code associated with data analysis performed using Python is available in reference [15].

To determine the illumination uniformity of 365 nm and 280 nm light, images of the fluorescent slide were opened in Fiji [16]. For each image, a line profile with a width of 50 pixels was taken horizontally across the field of view, and the intensity plotted as a function of distance. The standard deviation of the mean intensity was calculated for each illumination wavelength.

To analyse the relative fluorescence signal intensity of the QDs with excitation at 365 nm and 280 nm, a background corrected image pair with 365 nm and 280 nm excitation wavelengths was acquired by subtracting the average background intensity as measured in section 2.4. A binary mask of the 280 nm image was created using the Otsu thresholding algorithm [17]. This binary mask was applied to both the 365 nm and 280 nm QD images to isolate the regions of interest containing fluorescent signal from QDs in both images. 280nm:365nm intensity signal ratios were then calculated on a pixel-by pixel basis. Pixel locations in the masked 365 nm image with background-corrected intensities of 0 were identified and then discarded from both images to prevent infinite intensity ratios. Fluorescence intensity distributions from images excited with 280 nm and 365 nm were subjected to Welch’s t-test [18] under the null hypothesis that the intensity distributions have identical mean values, implying that there is no significant difference in emission intensity between excitation wavelengths.

To measure the photobleaching rate of QDs, a thresholding operation was performed using Fiji [16] and a binary mask of the cells was created. This was applied to the corresponding time-lapse image stack and the mean intensity of the cells were measured for each frame.

## 3. Results

When comparing the illumination uniformity of 365 nm and 280 nm illumination, it was found that 365 nm illumination varied by*±*4.98% of the mean across the field of view and the intensity of the 280 nm illumination varied by *±*3.23%.

Images of QD525-labelled HeLa cells excited with 280 nm and 365 nm light are shown in Figure 2. During acquisition of these images, imaging conditions such as camera exposure time and optical power at the specimen plane remained identical, with only the wavelength of excitation light changing between images. Figure 2(A) shows QD525-labelled cells excited with 365 nm light, and 2(B) shows cells excited with 280 nm light, with equal contrast adjustment. QD labelled cells appear visually brighter when excited with 280 nm light. This enhancement in fluorescence intensity, combined with minimally increased background intensity, also results in enhanced image signal-to-background.

**Figure 2:**
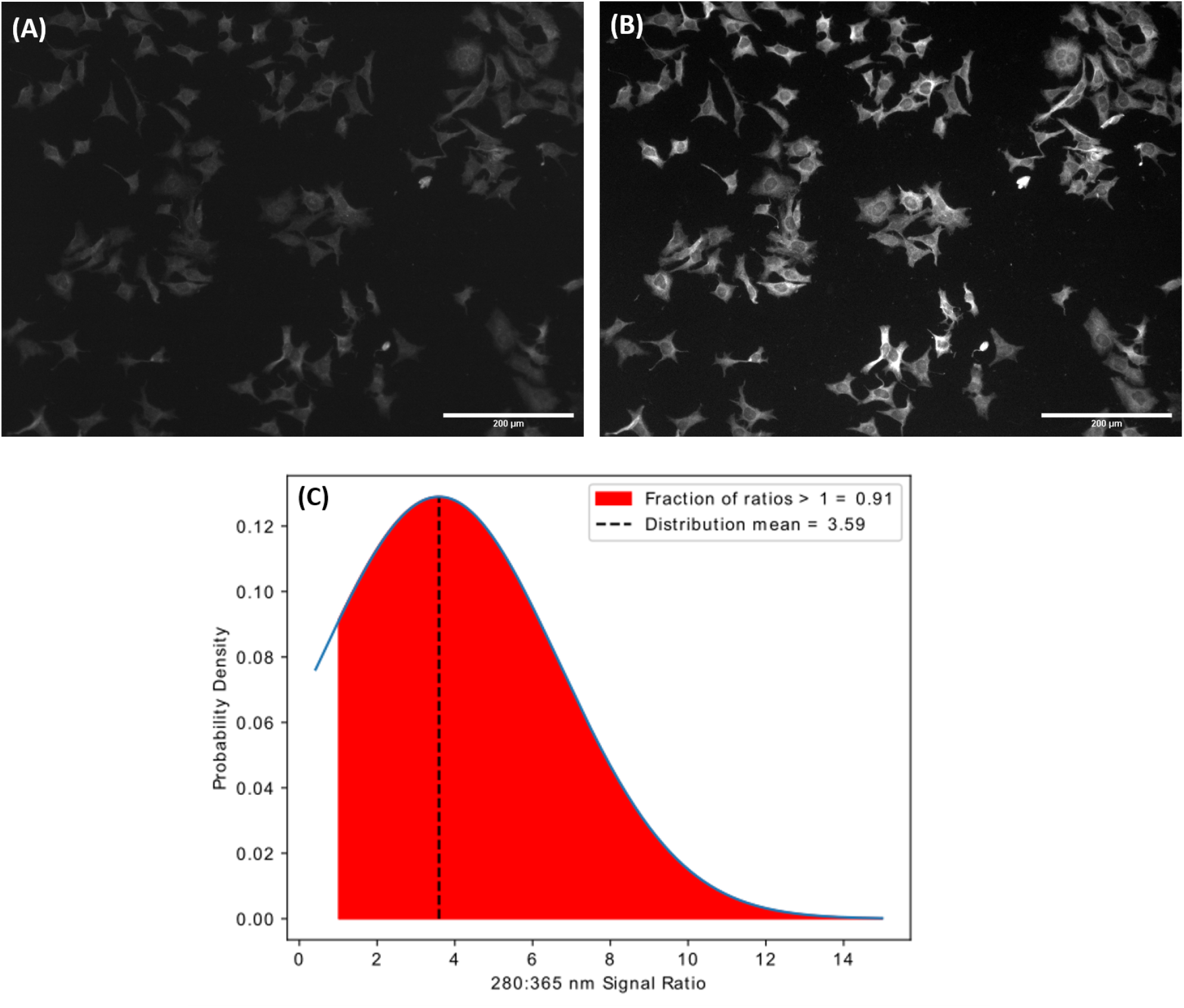
HeLa cells with QD525-labelled microtubules excited with (A) a wavelength of 365 nm and (B) a wavelength of 280 nm. (C) The probability distribution of the ratios of fluorescent signal from 280 nm and 365 nm excitation. The mean of the distribution is 3.59. The area shaded in red represents the the ratios > 1, which comprise 0.91 of the total cumulative probability density above 0.

To quantify the increase in fluorescence intensity from QDs excited with 280 nm light, a threshold of each image was performed using the Otsu method, and this threshold was applied to extract cellular regions of interest (ROIs) in each image as described in Section 2.5[17]. The intensity of each pixel within ROIs of the 280 nm excitation images were then compared with the pixel at the same location in the 365 nm excitation images, and a ratio of the fluorescent signal at 280 nm compared to 365 nm excitation was calculated for each pixel in the cellular ROIs. Figure 2(C) shows the distribution of 280nm:365nm ratios. The mean 280nm:365nm intensity ratio is 3.59, hence, on average, the intensity of QD525-labelled cells excited with 280 nm is 3.59-fold that of those excited with 365 nm. Furthermore, a two-sided t-test of 280 nm and 365 nm intensity distributions was performed under the null hypothesis that the distributions have identical mean values. This test yields a t-statistic of 472.43 and a p value of *≤*0.00001, confirming that the difference in mean fluorescence intensities of QDs excited with each wavelength is statistically significant.

To verify that this increase in excitation efficiency is applicable to multiple sizes of semiconductor QDs, imaging was repeated with HeLa cells with microtubules labelled using commercial QD605 streptavidin conjugates. HeLa cells with microtubules labelled using QD605 are shown in Figure 3, again excited with 365 nm light (A) and 280 nm light (B) and detected at wavelengths above 561 nm. Again, imaging conditions such as camera exposure time and optical power at the specimen plane remain identical for both images, with only the excitation wavelength changing. Data analysis was performed as before and the resulting distribution of QD intensity ratios can be found in Figure 3(C). Whilst we do see an increase in intensity associated with 280 nm excitation, this does not appear to be as pronounced as in the case of QD525-labelled cells. The mean 280nm:365nm intensity ratio is 2.03, hence, on average, the intensity of QD605-labelled cells excited with 280 nm is 2.03-fold that of those excited with 365 nm. Again, a two-sided t-test of 280 nm and 365 nm intensity distributions was performed. This test yields a t-statistic of 269.961 and a p-value *≤* 0.00001, confirming that the difference in mean fluorescence intensities of QDs excited with each wavelength is statistically significant.

**Figure 3:**
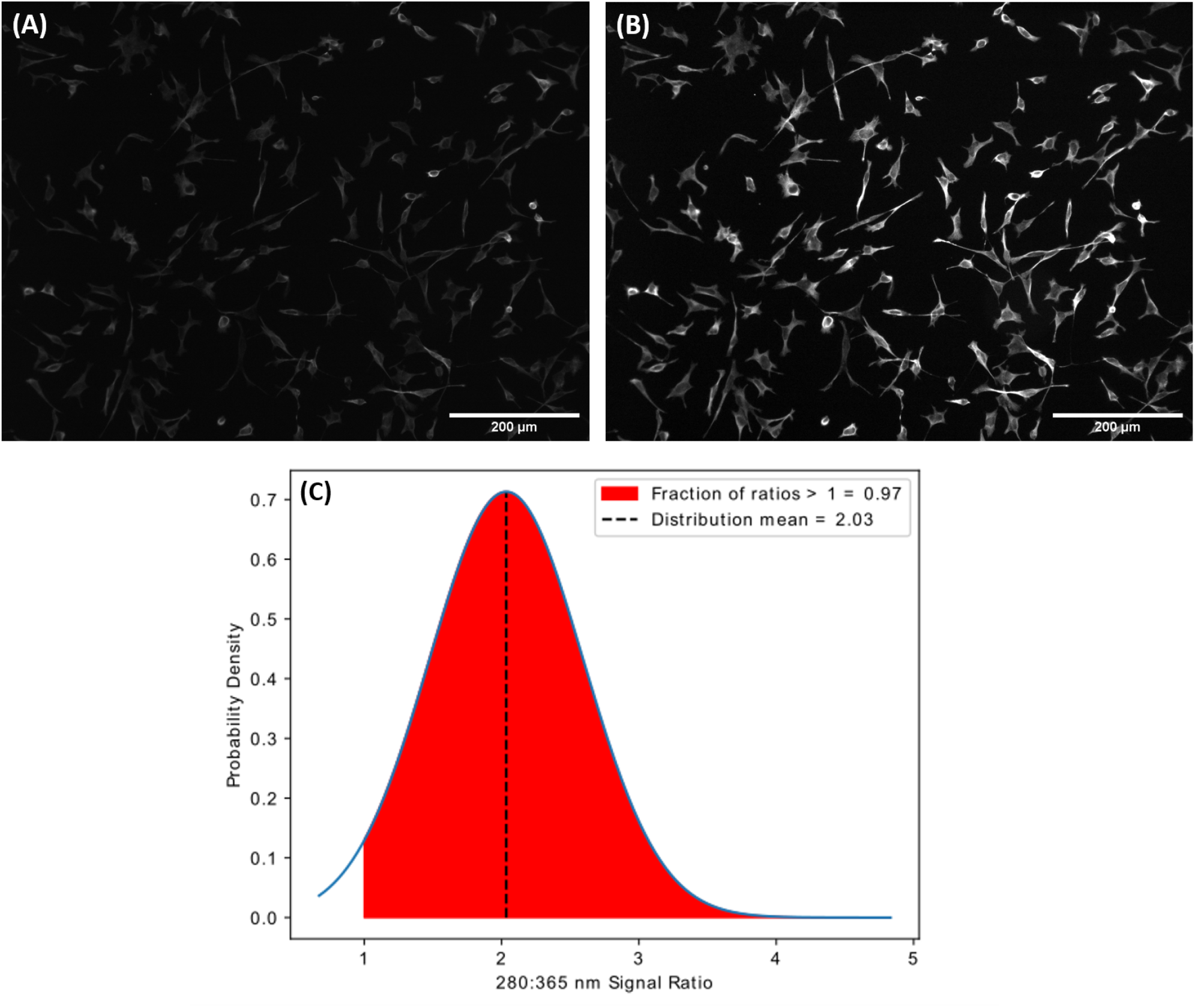
HeLa cells with QD605-labelled microtubules excited with (A) a wavelength of 365 nm and (B) a wavelength of 280 nm. (C) The probability distribution of the ratios of fluorescent signal from 280 nm and 365 nm excitation. The mean of the distribution is 2.03. The area shaded in red represents the the ratios > 1, which comprise 0.97 of the total cumulative probability density above 0.

The rate of photobleaching of QD605-labelled cells was investigated with both excitation wavelengths to ensure that the higher energy associated with 280 nm excitation did not have a more profound effect on photobleaching than 365 nm. After irradiating QD labelled cells each with 365 nm and 280 nm light for an 8-hour period, no evidence was found that 280 nm excitation causes increased photobleaching in commercial QDs when compared to 365 nm excitation (Figure 4). Fluorescence intensity from QDs was found to vary by 0.59% over the 8-hour period when irradiated with 365 nm light, and increase by 1.64% when irradiated with 280 nm light.

**Figure 4:**
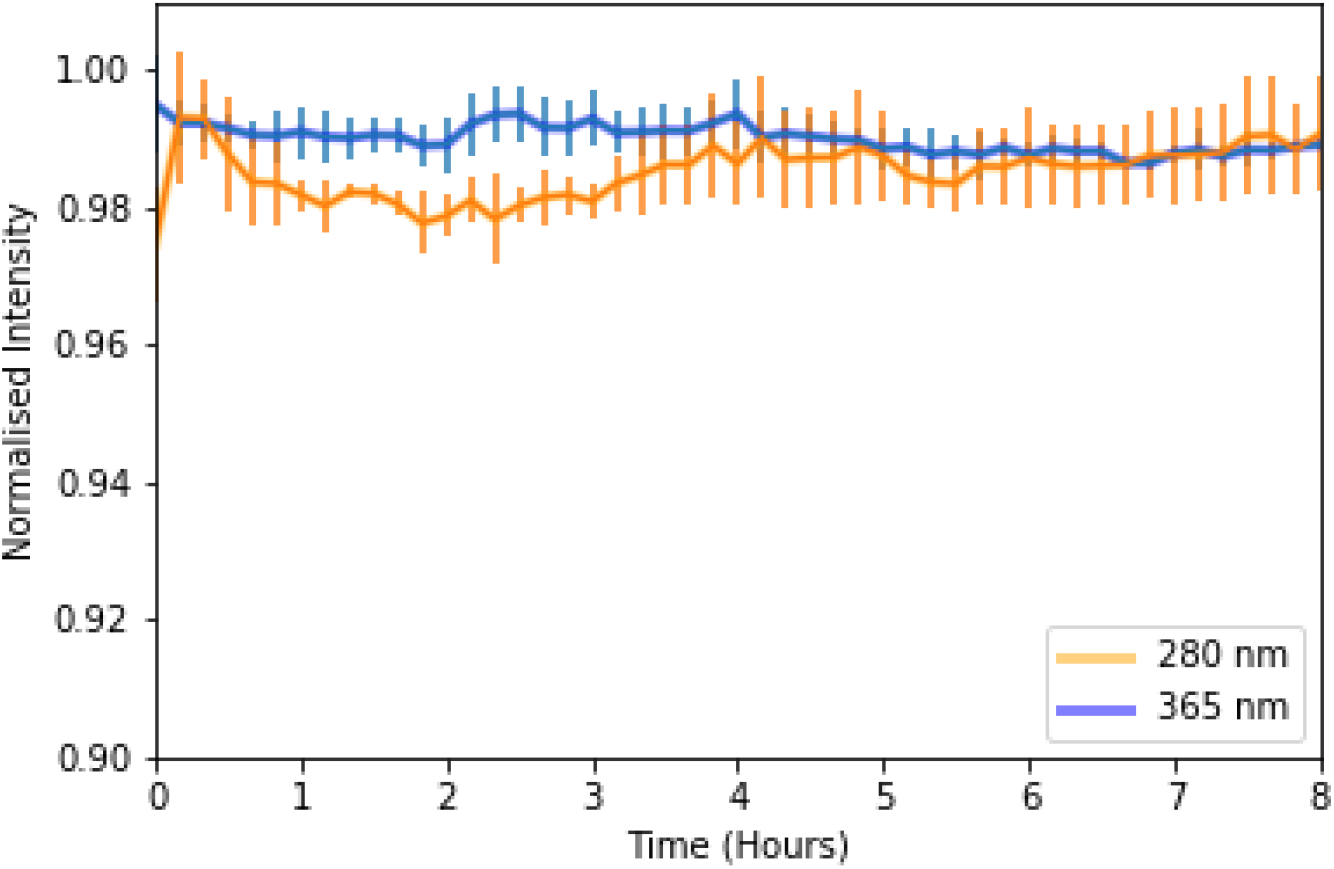
Mean intensity of QD605-labelled HeLa cells over an 8 hour period of constant irradiation with 365 nm (blue) and 280 nm (orange). Error bars represent the standard deviation of the mean.

## 4. Discussion

The observed increase in fluorescence intensity of QD-labelled cells is expected to apply to all sizes of commercial semiconductor QDs since all sizes of QDs have absorption spectra with broadly similar distributions [10]. Therefore, all sizes of QDs are likely to have a higher absorption efficiency at 280nm compared to longer wavelengths. However, although the absorption spectra distributions are similar, they are not identical for all QDs and the extent of the increase in fluorescence signal depends on the specific difference in absorption efficiency between 280nm and longer wavelengths for different sizes of QD.

Broad distributions in intensity ratios can partly be attributed to inhomogeneity of the illumination light. Whilst both illumination sources were aligned to achieve the best possible homogeneity of illumination across the field of view, it was not always possible to achieve this perfectly. As reported, some variations in intensity across the field of view occurred for both illumination wavelengths, affecting the mean increase in fluorescence intensity achieved using 280 nm excitation. In addition to this, while the size and absorption/emission properties of semiconductor QDs can be controlled via synthesis, not all synthesis methods result in QDs of one single size [19]. Therefore, within a sample of commercial QDs there will be a size tolerance leading to some variation in emission and absorption spectra [20, 21] which, as the increase in excitation efficiency is dependent on the shape of the absorption spectrum, can affect the mean increase in intensity between excitation wavelengths. Despite the substantial overlap in standard deviations from the mean fluorescence intensity at 280nm and 365nm excitation for both the QD525 and QD605 datasets, this does not correspond to significant instances where 365 nm excitation yields equivalent or brighter emission intensity. Indeed, the percentage of pixels where the 280nm:365nm intensity ratio is *>*1 (i.e. the percentage of pixels where the pixel in the 280nm excitation image has a higher intensity then the same pixel in the 365nm excitation image) is 98.88% and 99.28% for QD525 labelled cells and QD605 labelled cells, respectively. This near universal increase in intensity in favour of 280 nm excitation, coupled with p-values close to zero, confirms that these standard deviations from the mean do not detract from the conclusion that excitation of QDs with 280 nm light yields an increased fluorescence intensity.

Whilst using oblique illumination methods such as MUSE and others [22] can be advantageous in bypassing the need for quartz objectives, these techniques are currently limited to low magnification, long working distance lenses in order for light to bypass the objective at an angle and illuminate the specimen. In this regard, transmission illumination allows for much greater flexibility in objective lenses, including high magnification, high numerical aperture lenses which allow for more detailed imaging of cell specimens with improved resolution. Aside from MUSE, several other deep-UV microscopy techniques have relied on specialised objective lenses to image in either epifluorescence or brightfield modes [5, 23, 24, 25, 26]. These objectives, such as quartz or reflective objectives, are rarer than typical glass objectives, often more expensive and come in a very limited range of magnifications and numerical apertures. In addition to this, further modification to a commercial epifluorescence system would be required for UV transmission, including the replacement of internal lenses with quartz. The transmission fluorescence setup described here uses off-the-shelf optical components and does not involve any modification to the commercial microscope outside of the removal of the condenser lens, improving accessibility.

Although the fluorescence intensity of CdSe QDs does not show long-term degradation in when dispersed in an organic solution [27], increase in fluorescence intensity from QDs over time has been reported previously [28]. This has been attributed to carriers being transferred to surface traps present at the interface of the CdSe core and ZnS shell of the QDs or photo-assisted release of trapped carriers on the QD surface [27]. Further to this, there is also the possibility that the use of high-energy ultraviolet light could affect the thermal state of the specimen, e.g. heating of the gelvatol mounting medium, causing fluctuations in fluorescence intensity. This problem could be minimised by using aqueous mountant such as in live-cell imaging experiments. Since minimal photobleaching of QDs occurs in the short term under either excitation wavelength, this makes it unlikely that the intensity difference observed in QDs excited by 280 nm vs 365 nm light is caused by photobleaching. In addition, the images with 365 nm excitation were acquired before images with 280 nm excitation, further ruling out the possibility of photobleaching affecting intensity ratios as the second image acquired is always brighter than the first.

To build on this work, we plan to investigate whether the observed increase in fluorescence from QDs excited with 280 nm light can be applied to study live cells. Whilst 280 nm light is known to be damaging to cells due to its proximity to the absorption peak of DNA (□260 nm) [29], previous studies using 280 nm light in live-cell imaging have proven successful, with authors able to image cells for 6 hours before the onset of cell death [5] by triggering the excitation light, limiting exposure of live cells to 280 nm light. It is hoped that by using this approach, we can exploit the high fluorescence intensity associated with 280 nm excitation of QDs to study cellular dynamics such as migration and mitosis with minimal UV-induced toxicity.

## 5. Conclusion

We have demonstrated a significant improvement in fluorescence intensity of semiconductor QDs within the cellular environment when using 280 nm excitation. We report up to a 3.59-fold increase in fluorescence intensity when using 280 nm excitation when compared to 365 nm excitation, resulting in significantly enhanced image quality. This increase is expected to apply to all emission varieties of commercial semiconductor QDs due to their common absorption spectra. In addition to this, we find no significant increase in photobleaching of QDs when illuminated with 280 nm light over an 8-hour period when compared to 365 nm light. We anticipate that by minimising UV exposure to live cells, the high fluorescence intensity associated with 280 nm excitation of QDs can, in future, be exploited in live-cell studies with minimal UV-induced toxicity.

## Acknowledgments

This work was funded by Medical Research Scotland (PhD-1157-2017) and CoolLED Ltd.NH and GMcC are part-funded by the Biotechnology and Biological Sciences Research Council (BB/T011602/1). The authors would like to thank Gerard Whoriskey, Alex Gramann and Luther Hindley at CoolLED Ltd for assisting with the 280 nm LED used in this paper and Lisa Kölln at University of Strathclyde for useful discussions relating to antibody labelling.

## Data Availability Statement

The data generated and analysed during this study is available in the repository in reference [15].

